# Vascular Transcription Factor TMO5 Regulates PIN1 Polarity and Organogenesis Downstream of MONOPTEROS in the Shoot Meristem

**DOI:** 10.1101/2024.11.09.622801

**Authors:** Abdul Kareem, Carolyn Ohno, Marcus G. Heisler

## Abstract

Plants continuously produce lateral organs, such as leaves and flowers, from the shoot apical meristem (SAM). This process is guided by the accumulation of the plant hormone auxin and the polar localization of the efflux protein PIN-FORMED1 (PIN1). The transcription factor MONOPTEROS (MP) plays a crucial role in orienting PIN1 polarity, thereby facilitating auxin-driven organogenesis. In this study, we investigated how MP regulates PIN1 polarity and organogenesis, discovering that the downstream vascular transcription factor TMO5 regulates PIN1 polarity non-cell-autonomously and promotes organ initiation in the SAM. Furthermore, we find that exogenous auxin and cytokinin can partially rescue the organ defects associated with loss of MP and TMO5 activity. Collectively, these findings uncover a novel role for TMO5 in controlling PIN1 polarity and driving organogenesis by coordinating hormonal signalling.

## Introduction

Plants generate lateral organs such as leaves and flowers periodically from the periphery of the shoot apex or shoot apical meristem (SAM). While the resulting patterns, or phyllotactic arrangements of organs, have fascinated biologists and artists alike for hundreds of years (Jean and Barab, 1998), our understanding of the underlying molecular mechanisms remains far from complete. Organ outgrowth is triggered by local accumulation of the plant hormone auxin, which is transported to specific locations within the SAM periphery via the polarly localised membrane-bound efflux carrier protein PIN-FORMED1 (PIN1) (Heisler et al., 2005; Okada et al., 1991; Reinhardt et al., 2000; Reinhardt et al., 2003). Studies have now demonstrated that the patterns of PIN1 polarity required to concentrate auxin at organ initiation sites occur due to positive feedback between auxin signalling and the polarity of PIN1 localization in neighbouring cells (Bhatia et al., 2016). This feedback loop is mediated by the MONOPTEROS (MP) or AUXIN RESPONSE FACTOR 5 (ARF5) transcription factor (Hardtke and Berleth, 1998; Przemeck et al., 1996), which orients PIN1 polarity non-cell autonomously towards MP-expressing cells (Bhatia et al., 2016). Understanding how MP activity results in changes to PIN1 polarity remains a central question in the field.

Previous work has identified several downstream targets of MP (Cole et al., 2009; Konishi et al., 2015; Schlereth et al., 2010; Yamaguchi et al., 2013). Some of these targets, including LEAFY (LFY), AINTEGUMENTA (ANT), and AINTEGUMENTA-LIKE6 (AIL6), are known to contribute to flower development. However, the degree of rescue of the flowerless *mp* phenotype by LFY and ANT expression is limited (Yamaguchi et al., 2013). Other known MP targets, including the Target of MONOPTEROS 5 (TMO5) and Target of MONOPTEROS 6 (TMO6) genes act downstream of MP in vascular tissue and embryonic root development (Schlereth et al., 2010). For instance, the bHLH transcription factor TMO5 promotes periclinal divisions in developing root vasculature through a cytokinin biosynthesis pathway (De Rybel et al., 2014; De Rybel et al., 2013; Ohashi-Ito et al., 2014). When mis-expressed with its dimerization partner LONESOME HIGHWAY (LHW), TMO5 triggers ectopic leaf initiation on the petiole (De Rybel et al., 2014) suggesting a more general role as a promoter of growth and organ outgrowth in shoots. Furthermore, a recent study has shown that TMO5 and its homologues, TMO5-LIKE2 (T5L2) and T5L3 are expressed in the SAM (Mor et al., 2022).

Here, we investigate the role of several MP target genes in lateral organ formation. We demonstrate that like MP, TMO5 can regulate PIN1 polarity in the SAM epidermis non-cell autonomously and promote organogenesis. We also present evidence that TMO5 and its close homologues promote organogenesis downstream of MP by amplifying and triggering auxin and cytokinin signalling respectively.

## Results and Discussion

To identify genes downstream of MP involved in regulating organ initiation and PIN1 polarity in the shoot, we investigated several known candidate genes including TMO5 (De Rybel et al., 2014; De Rybel et al., 2013), TMO6 (Miyashima et al., 2019; Smet et al., 2019) and DOF5.8 (Konishi et al., 2015) using translational fusions to yellow fluorescent protein for energy transfer (YPET), including TMO5::TMO5-2YPET, TMO6::TMO6-2YPET and DOF5.8::DOF5.8-YPET. In both vegetative and inflorescence meristems, expression of both TMO5::TMO5-2YPET and DOF5.8::DOF5.8-YPET was detected in epidermal and sub-epidermal layers of organ primordia in a pattern partly overlapped with PIN1 expression (Fig. 1A,B, Fig. S1A,B,D). In contrast, TMO6::TMO6-2YPET expression was only detected in sub-epidermal and inner cells of organ primordia (Fig. 1C, Fig. S1F). To test whether the expression of these genes depends on MP, we analysed our reporters in *mp* mutant meristems. We found almost no detectable TMO5 expression in the strong *mp* mutant (*mpT370*) dome meristem or pin-like inflorescence meristem (Fig. 1D). However, TMO5 expression was detected in the rarely formed *mp* mutant leaves (Fig. S1C). Similarly, DOF5.8::DOF5.8-YPET expression was low in the *mp* pin-like inflorescence meristem but was detected in leaf primordia that had developed from the vegetative meristem (Fig. 1E, Fig. S1E). TMO6::TMO6-2YPET expression was still detected in the *mpT370* mutant pin-like meristem but this was reduced compared to the wild type (Fig. 1F, S1G). These data confirm that TMO5, DOF5.8 and TMO6 expression correlates with organogenesis and that the extent of their expression depends on MP activity.

**Fig. 1.**
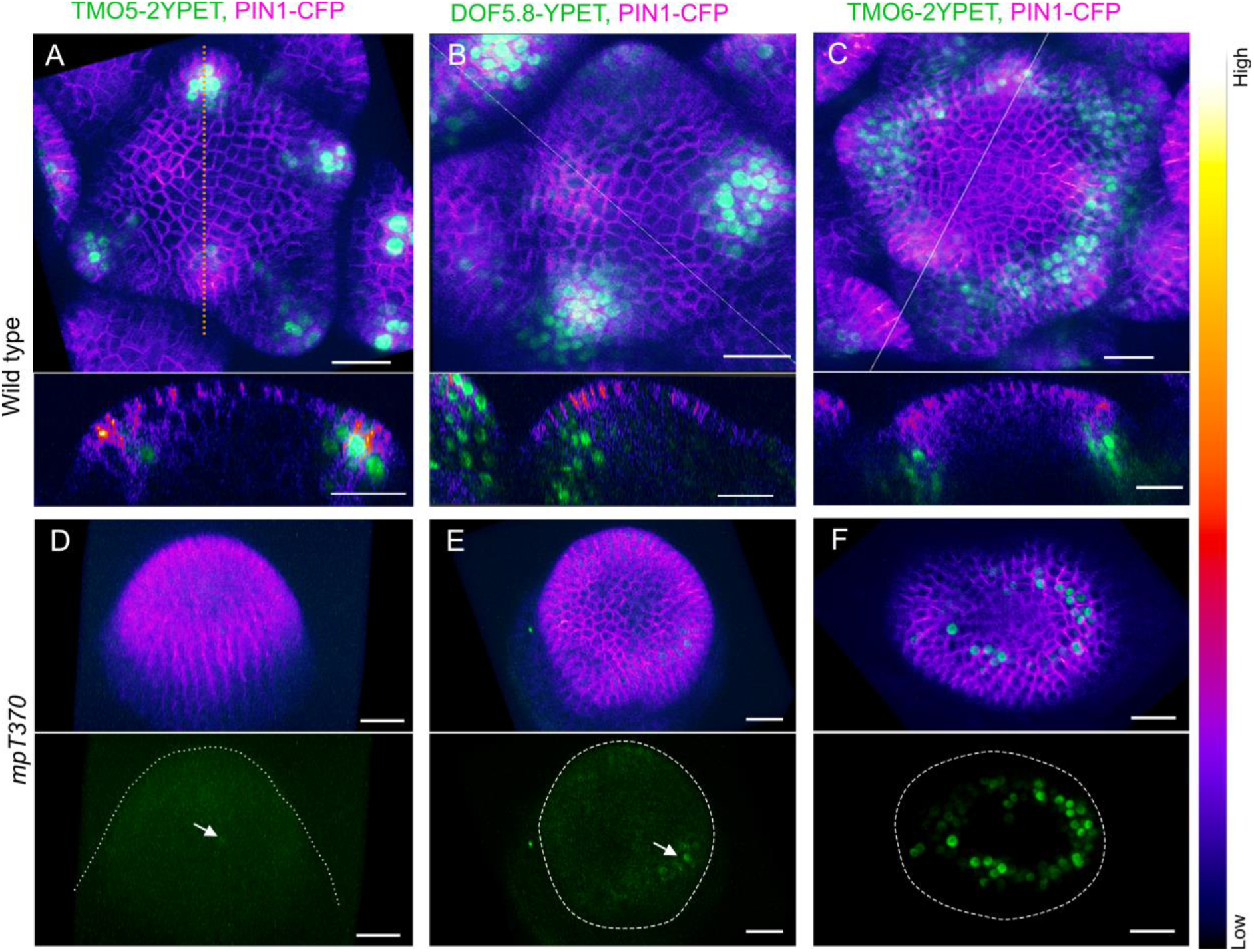
Expression patterns of TMO5, DOF5.8, and TMO6 in wild type and *mp* mutant meristems. (A) TMO5::TMO5-2YPET expression (green) in wild type inflorescence meristem, partially overlaps with PIN1::PIN1-CFP expression (magenta). (B) DOF5.8::DOF5.8-YPETexpression (green) in wild type inflorescence meristem, overlaps with PIN1::PIN1-CFP signal. (C) TMO6::TMO6-2YPET expression (green) in wild type inflorescence meristem. Lower panels in (A–C) show corresponding longitudinal optical sections. (D–F) TMO5, DOF5.8, and TMO6 expression in the *mp-*T370 mutant dome meristem, with lower panels displaying the green channel. Dotted lines indicate meristem boundaries. Barely detectable TMO5 and DOF5.8 signals are marked with arrows. Scale bar = 20 μm.

We next tested whether localized expression of these genes in the *mp* mutant meristem could promote organ formation and alter PIN1 localisation patterns. Towards this end, we generated small clones of cells expressing these genes individually in the *mp* mutant (*mp*T370 and *mp*B4149) shoot meristem, harbouring a PIN1-CFP (or PIN1-GFP) marker. This was accomplished using an inducible Cre-lox recombination-mediated genetic mosaic system targeted to the CLAVATA3 (CLV3) expression domain (pCLV3:CRE-GR+pUBQ10::GENE-2YPET), as described previously (Bhatia et al., 2016). We found the appearance of clones expressing TMO5, DOF5.8 or TMO6 in the *mp* dome-shaped shoot meristem within two days of DEX induction of Cre-GR. By 3-5 days, PIN1 expression levels increased within the clones expressing TMO5 but not DOF5.8, nor TMO6 (Fig. S2A-C, E-H). Clonal induction of TMO5, but not DOF5.8 or TMO6, also led to organ outgrowth within 6-7 days of induction, resulting in the formation of flower-like structures (Fig. 2A-E, F-J Fig. S2D). Transient ubiquitous expression of TMO5 also induced flower formation in *mp* mutants (Fig. S2I), while local clonal activation in wild type plants resulted in altered organ positioning (Fig. 2K,L). Examining TMO5 clones more closely, we found that similar to MP (Bhatia et al., 2016), localized TMO5 activity was sufficient to induce a localized PIN1 polarity convergence (Fig. 2M,N), with PIN1 localization becoming polarized towards the TMO5 clones in cells adjacent to the clones (Fig. 2O,P, S2I,J). This demonstrates that TMO5 can influence PIN1 polarity in a non-cell-autonomous manner. Additionally, we observed activation of PIN1 expression in sub-epidermal cells located in the L2 (layer 2) in response to overlying TMO5 clones in the epidermis (Fig. 2P,Q). We also detected upregulation of PIN1 expression and convergence in the epidermal cells in response to underlying TMO5 clones in sub-epidermal L2 cells (Fig. 2Q, S2K). These sub-epidermal TMO5 clones were sufficient to induce organ outgrowth in the *mp* pin-like meristem, similar to the effect of epidermal clones (Fig. 2P,Q, S2K). Altogether, these data support the proposal that TMO5, but not TMO6 or DOF5.8, act downstream of MP to promote organ outgrowth and polarize PIN1 non-cell autonomously.

**Fig. 2.**
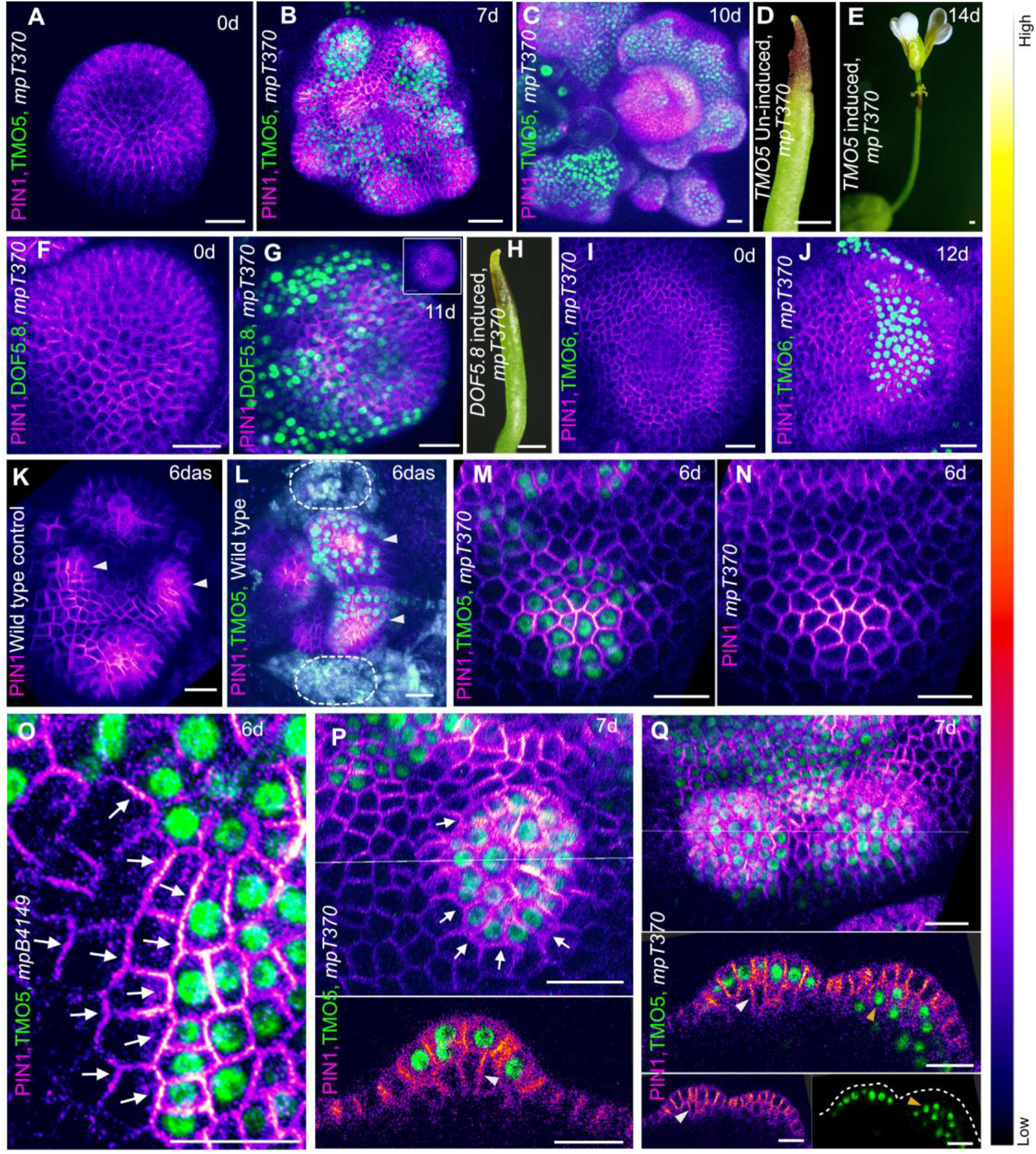
TMO5 promotes PIN1 polarity and organogenesis independent of MP. (A-C) Activation of PIN1-CFP expression (magenta), convergence formation, and organ outgrowth in *mp* mutant shoot meristems after clonal activation of TMO5-2YPET (green). Panels (A) and (B) show the same meristem over time, while panel (C) presents a representative meristem. (D,E) Flower formation on *mp* pin-like inflorescence meristem following clonal activation of TMO5-2YPET. (F-J) No effect on PIN1 localization or organ outgrowth in *mp* mutant meristems with local activation of DOF5.8-YPET (F-H) or TMO6-YPET (I,J). Inset in (G) shows the meristem with the PIN1-CFP channel alone for clearer visualization. (K) Six-day-old wild type vegetative meristem with PIN1-GFP expression, showing four leaves. Arrowheads indicate the third and fourth leaves. (L) Six-day-old wild type vegetative meristem with PIN1-GFP expression, showing altered positioning of the third and fourth leaves (arrowheads) due to clonal activation of TMO5-2YPET (green). The first and second leaves have been removed (dotted circles) to visualize the altered organ positioning. (M, N) Localized PIN1-CFP convergence formation in TMO5-expressing clones in the *mp* mutant meristem, with a separate PIN1-CFP channel for clearer visualization in (N). (O) Polarization of PIN1-GFP towards TMO5-expressing clones in neighboring cells. (P, Q) Activation of PIN1 in sub-epidermal cells (white arrowhead) due to TMO5 clones in the overlying epidermis, and upregulation of PIN1 expression and convergence in epidermal cells due to underlying sub-epidermal TMO5 clones (orange arrowhead), leading to organ outgrowth. Lower panels in (P, Q) show longitudinal optical sections corresponding to the lines in the upper panels. Dotted lines in (Q) mark organ boundary. Arrows indicate PIN1 polarity. Scale bar = 20 µm, except for (D, E, and H), where it is 1 mm.

To determine whether TMO5 contributes to organ formation during normal development, we analysed organogenesis in *tmo5* single mutants but found no defects in organ initiation (Fig. S3A) (De Rybel et al., 2013; Vera-Sirera et al., 2015). Therefore, we investigated the function of genes closely related to TMO5 including TMO5-LIKE1 (T5L1) T5L2, T5L3 (De Rybel et al., 2013), as well as a newly identified homologue, AT2G40200, which we named T5L4 (Fig. S3A). Like other homologs, AT2G40200/T5L4 has been reported to dimerize with the partnering protein LHW in vivo (De Rybel et al., 2013). We detected the expression of T5L1, T5L2, and T5L3 in the shoot apical meristem, consistent with previous report (Mor et al., 2022), and found that T5L4 was also expressed in the meristem (Fig. 3A-D). Since single mutants *t5l1, t5l2, t5l3*, and *t5l4* did not display any defects in organ (De Rybel et al., 2013; Vera-Sirera et al., 2015) (Fig. S3B), we investigated higher order mutants. While the *tmo5 t5l2 t5l3* triple mutant displayed a significant reduction in the number of flowers formed in the inflorescence compared to wild type, the *tmo5 t5l4* double and *tmo5 t5l1 t5l4* triple mutants showed a greater reduction in the number of flowers formed (Fig. 3E). The *tmo5 t5l1 t5l2 t5l3* quad mutant showed the most severe phenotype and produced no flowers (Fig. 3E-G). Though the quad mutant germinated similar to the wild type, it displayed retarded root and shoot development and a tiny stature (Fig. 3G, S3C,D). Shoot growth terminated after the 6-8 rosette leaf stage without producing an inflorescence (Fig. 3G). We next analysed the phyllotaxis of *tmo5 t5l1 t5l4* and *tmo5 t5l2 t5l3* triple mutants and observed an abnormal pattern of siliques (Fig. 3H) indicating that TMO5 and T5Ls contribute to spatial organ positioning.

**Fig. 3.**
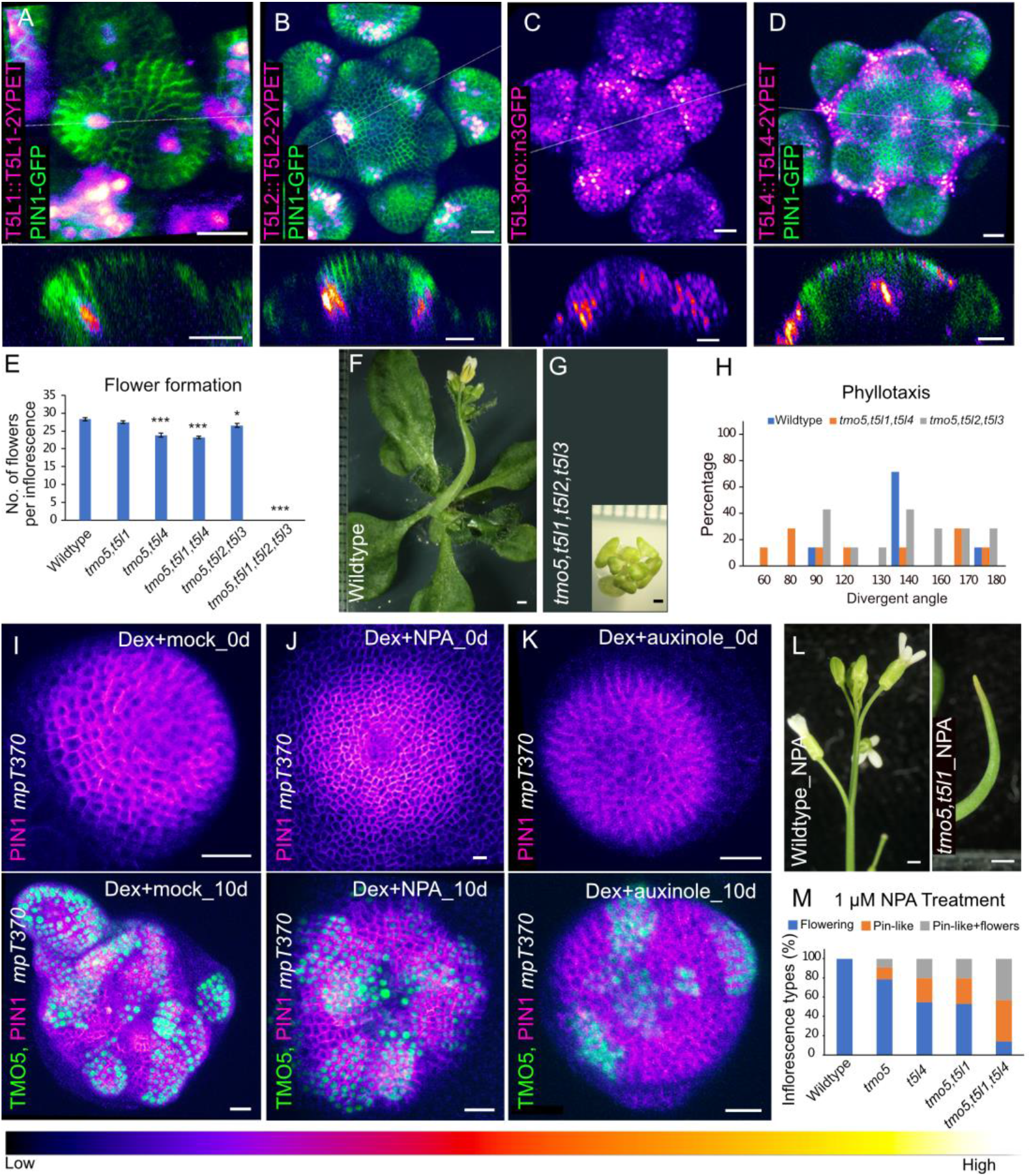
Role of TMO5 and related T5L genes in organ formation. (A-D) Expression patterns of T5L1, T5L2, T5L3, and T5L4 reporters (magenta) in the wild type inflorescence meristem with or without PIN1-GFP (green). Lower panels show longitudinal optical sections corresponding to the lines in the upper panels. (E) Average number of flowers formed per inflorescence in wild type and *tmo5/t5l* mutants. (n= 20 plants per genotype; mean±SEM; *p<0.5; ***p<0.001; Student’s t-test). (F,G) Growth phenotypes of the homozygous *tmo5* quad (*tmo5,t5l1,t5l2,t5l3*) mutant in comparison to wild type. (H) Phyllotaxy phenotypes in *tmo5, t5l1, t5l4* and *tmo5, t5l2, t5l3* triple mutants, measured by the divergent angle and compared to the wild type, which has a divergent angle of 137°. (n= 7-14 plants). (I) Organ formation in mock treated *mp* mutant meristem following TMO5-2YPET (green) clonal activation. Upper panel: 0 days post induction. Lower: 10 days post induction. (J) Organ formation in 10 mM NPA treated *mp* mutant meristem after TMO5-2YPET clonal activation. Upper panel: 0 days post induction. Lower: 10 days post induction. (K) Suppression of TMO5-mediated organ formation in *mp* mutant meristems treated with 100μM auxinole, an auxin signaling inhibitor. Upper panel: 0 days post induction. Lower: 10 days post induction. (L) Formation of pin-like inflorescence in *tmo5,t5l1* mutants, but not in wild type, after 1 μM NPA treatment. (M) Frequency of formation of inflorescence types (flowering, pin-like, and pin-like with flowers) in wild type and *tmo5/t5l* mutants following treatment with 1 μM NPA. (n=15-35 plants per genotype). Scale bars: 1 mm, except in (I-K) where it is 20 μm.

Given that T5L1 regulates the auxin biosynthesis gene *YUCCA4* (*YUC4*) during vascular development in the root (Ohashi-Ito et al., 2019), we assessed the organogenic activity of TMO5 clones in the *mp* mutant to understand how TMO5 promotes organ formation in relation to auxin. We found that clonal TMO5 induction was still able to induce organ formation in the presence of 10μM NPA, a concentration of NPA sufficient to cause a pin-like inflorescence in wild type and suppress organ outgrowth in the *mp* mutant (Reinhardt et al., 2000; Schuetz et al., 2008), albeit with delayed and reduced frequency (Fig. 3 I,J). However, when auxin signalling was reduced by treating with 100uM auxinole, an inhibitor of auxin signalling, TMO5-induced organ formation was completely suppressed (Fig. 3K), indicating that TMO5 requires residual auxin signalling in the *mp* mutant to induce organ formation. To test whether TMO5 function is also sensitive to auxin in a wild type context, we treated *tmo5* mutant combinations with upon 1μM NPA treatment, a concentration that is usually insufficient to elicit organogenesis defects in wild type (Fig. 3L,M). Under such conditions we found that *tmo5* single mutants produced a leafless meristem structure after making the first pair of leaves (Fig. S3E,F) while later, a pin-like inflorescence developed (Fig. S3G). The *t5l4* single, *tmo5,t5l1* and *tmo5,t5l4* double and *tmo5,t5l1,t5l4* triple mutants also produced pin-like inflorescences upon 1μM NPA treatment (Fig. 3L,M, S3H-L) revealing that TMO5 and T5L genes normally act in synergy with auxin and its polar transport to control organ formation.

Besides auxin, TMO5-LHW heterodimers are known to regulate cytokinin biosynthesis and signalling to induce periclinal cell division during vascular development (De Rybel et al., 2013; Ohashi-Ito et al., 2014). We therefore tested whether the promotion of organogenesis by TMO5 in the *mp* mutant meristem could be phenocopied by exogenous applications of auxin or cytokinin. Indeed, we found that external application of cytokinin could trigger growth from the *mp* vegetative meristem (Fig. 4A-C). However, cytokinin application alone did not initiate outgrowth from the *mp* pin-like inflorescence meristem. Instead, we found that a combined application of auxin and cytokinin triggered organ-like outgrowth from the *mp* pin-like inflorescence meristem (Fig. 4D). Next, we tested if auxin and cytokinin treatment can rescue the organ outgrowth defect of the *tmo5,t5l1,t5l2,t5l3* quadruple mutant shoot. We found that the one-off cytokinin application with 10µM trans-zeatin (TZ) at shoot tip promoted new leaf formation in the 5day old quadruple mutant and thus rescued the vegetative defect (Fig. 4E,G,H, S4A). In contrast, auxin application (1µM 1-Naphthaleneacetic acid, NAA) did not rescue leaf formation in the mutant (Fig. 4E,I, S4A) indicating that cytokinin is the primary limiting factor in the *tmo5,t5l1,t5l2,t5l3* mutant for leaf formation. Although a one-off cytokinin application also led to inflorescence formation, only a few flowers formed on the inflorescence (Fig. 4H). Therefore, we applied auxin to the cytokinin-pre-treated quad mutants to test if auxin is the limiting factor for flower formation. Indeed, we found that auxin treatment enhanced flower formation (Fig.4J). However, the sole application of auxin was insufficient to rescue leaf or flower formation in the quad. These data therefore indicate that cytokinin can substitute for the role of TMO5/T5L in leaf formation while both cytokinin and auxin are required to complement the role of TMO5/T5L in flower formation.

**Fig. 4.**
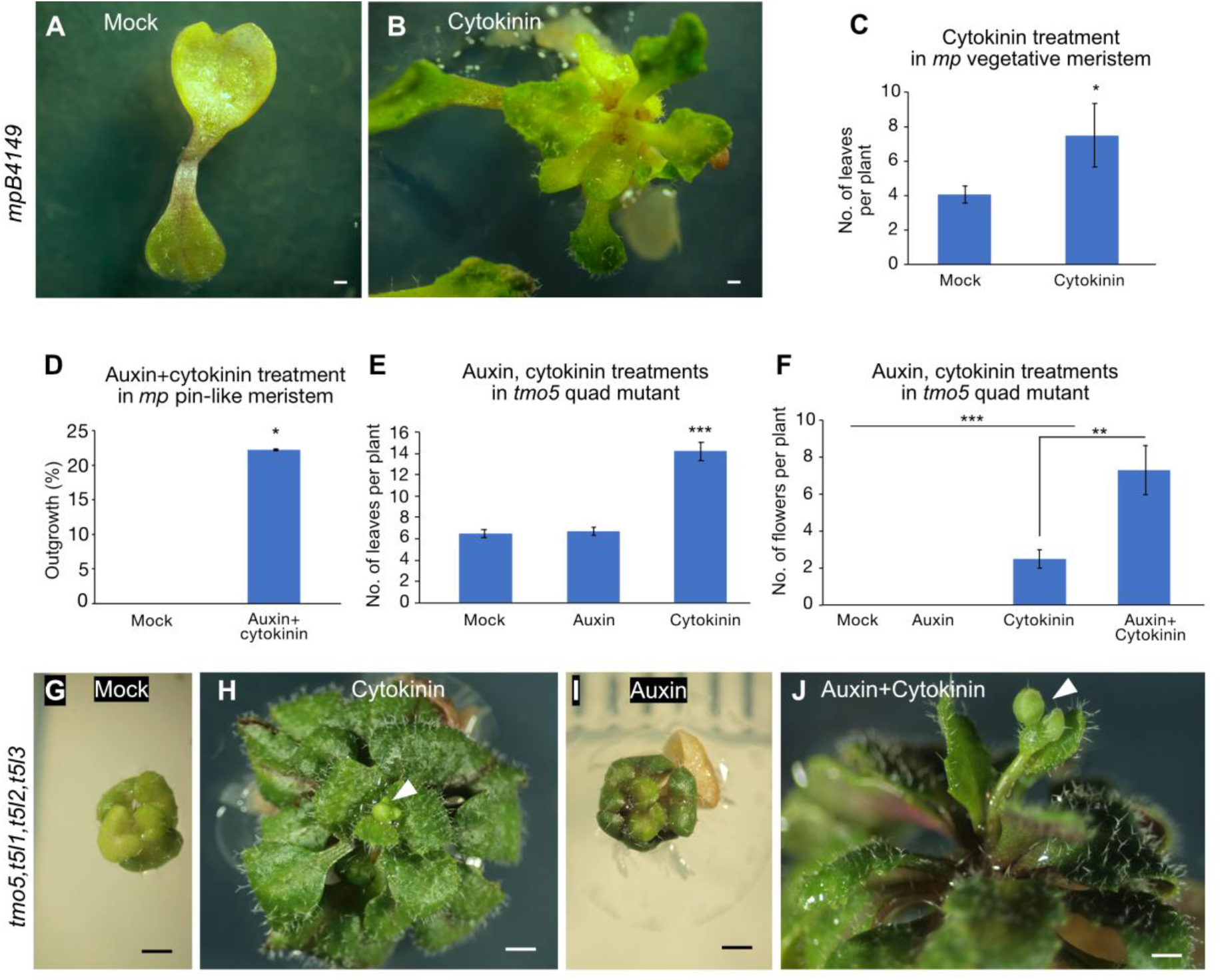
Cytokinin and Auxin Substitute TMO5/T5L Function for Organ Formation. (A,B) Leaf formation in *mp* mutant following 13 days of cytokinin treatment with 10 μM Trans-zeatin (TZ) (C) Average number of leaves per plant in the *mp* mutant following 10 μM Trans-zeatin (TZ) treatment. (n=10-5 plants per treatment; mean±SEM; *p<0.5; Student’s t-test). (D) Frequency of organ-like outgrowth in *mp* pin-like inflorescence meristem after treatment with both auxin (5 mM IAA) and cytokinin (1 mM TZ) treatmant. (n=18 plants per treatment; mean±SEM; *p<0.5; Student’s t-test). (E) Average number of leaves per plant in the *tmo5,t5l1,t5l2,t5l3* quad mutant following auxin (1uM NAA) and cytokinin (10 μM TZ) treatment. (n=10 plants per treatment; mean±SEM; ***p<0.001; Student’s t-test). (F) Average number of flowers formed per plant in *tmo5,t5l1,t5l2,t5l3* quadruple mutant after auxin (1uM NAA) and cytokinin (10 μM TZ) treatment. (n=10 plants per treatment; mean±SEM; **p<0.01; ***p<0.001; Student’s t-test). (G-I) Leaf and flower formation in the *tmo5,t5l1,t5l2,t5l3* quadruple mutant after cytokinin and auxin treatment. Single 10 μM TZ application rescues leaf formation and induces flowering (H); 1 μM NAA alone has no effect (I) but enhances flowering following cytokinin application (J). Arrowheads in (H and J) indicate flower buds. Scale bar = 1 mm.

Taken together, our data indicate that TMO5 regulates leaf outgrowth in the vegetative meristem by activating cytokinin while in the inflorescence meristem, TMO5 promotes flower formation by regulating both auxin and cytokinin. This scenario is consistent with the requirement for these hormones during organ formation in the dark when auxin transport is compromised (Yoshida et al., 2011) and is consistent with organ rescue previously observed in *mp amp1* double mutants, which have elevated cytokinin levels (Vidaurre et al., 2007). Given TMO5 is itself auxin-regulated (De Rybel et al., 2013) and predominantly expressed sub-epidermally (this study), our data also suggest that in the inflorescence meristem at least, organogenesis involves an amplification process starting from an auxin trigger that encompasses multiple cell layers. This not only includes the concentration of auxin locally via PIN1 mediated transport (Bhatia et al., 2016), but also, via the local induction of TMO5-mediated auxin synthesis or signalling. As it has been established that the cytokinin synthesis genes LOG3 and LOG4 are targets of LHW, which heterodimerizes with TMO5 and its homologs in the shoot (Mor et al., 2022), our work also suggests LOG3 and LOG4 contribute to flower formation in addition to promoting meristem function (Chickarmane et al., 2012). While T5L1 is known to promote auxin biosynthesis in roots (Ohashi-Ito et al., 2019), further work is required to better understand how TMO5 and its homologs feedback to auxin signalling and how auxin and cytokinin ultimately work in conjunction to promote organ outgrowth.

## Materials and Methods

### Plant materials and growth conditions

*Arabidopsis thaliana* ecotype Landsberg erecta (L*er*) or Columbia-0 (Col-0) was used as the wild type in this study. The *mp-*T370 mutant allele is in the L*er* ecotype and *mp-*B4149 allele is in the *Utrecht* ecotype (Bhatia et al., 2016; Hardtke and Berleth, 1998; Weijers et al., 2006). The remaining mutants used in this study including *tmo5, t5l1, t5l2* and *t5l3* (De Rybel et al., 2013) as well as *t5l4* (AT2G40200, SALK_008269.47.60, N508269) (this study) are in Col-0 background. The *t5l4* mutant was genotyped using the primers listed in Table S1. The *tmo5,t5l1* double and *tmo5,t5l1,t5l2,t5l3* quadruple mutants were described previously with genotyping details (De Rybel et al., 2013). The *tmo5,t5l2,t5l3* triple mutants was derived from *tmo5,t5l1,t5l2,t5l3* quadruple mutant siblings, where *t5l1* was segregating. The *tmo5,t5l4* double mutant was generated by crossing the respective single mutants and the *tmo5,t5l1,t5l4* triple mutant was produced by crossing *tmo5,t5l4* and *tmo5,t5l1* double mutants. All mutant combinations were genotyped using the primers listed in Table S1. The T5L3pro::n3GFP reporter was described previously (De Rybel et al., 2013). Other transgenic lines include *mp*-T370 mutants carrying the pPIN1::PIN1-CFP reporter, which were transformed with either TMO5::TMO5-2YPET, TMO6::TMO6-2YPET, DOF5.8::DOF5.8-YPET, or a genetic mosaic construct incorporating TMO5, TMO6, and DOF5.8 (details provided below). Additionally, the translational reporters T5L1::T5L1-2YPET, T5L2::T5L2-2YPET, or T5L4::T5L4-2YPET were introduced into wild type plants carrying pPIN1::PIN1-GFP.

Seeds were surface sterilized with 70% (v/v) ethanol for 10 minutes. The seeds were then put on a sterile filter paper to dry out and to remove ethanol content. The sterilized seeds were plated on 1X Murashige and Skoog (MS) basal salt mixture (Sigma M5524), 1% sucrose, 0.5g/L MES 2-(MN-morpholino)-ethane sulfonic acid (Sigma M2933), 0.8% Bacto Agar (BD) and 1% MS vitamins (Sigma M3900). pH was adjusted to 5.7 with 1M KOH. After 2 days of stratification at 4°C, the seed plates were moved to the growth room at 22°C under continuous light. For imaging of wild type inflorescence meristems, plants were grown on soil at 18C in short day conditions (16h/8h).

Construction of reporters and transgenic plants For genetic mosaic analyses using the CRE/Lox system, we generated stable transgenic lines containing a template for sectoring TMO5 (UBQ10p:lox spacer lox:GENE-2YPET) or DOF5.8 or TMO6 (UBQ10p:lox spacer lox:GENE-YPET) along with a dexamethasone-inducible CRE Recombinase (CLV3p:CRE-GR) (Bhatia et al., 2016). Here, ‘GENE’ refers to DOF5.8, TMO5, or TMO6. To create UBQ10p:lox spacer lox:GENE-2xYPET, we initially cloned a 4.6 kb SfiI-BamHI fragment from UBQ10p:lox spacer lox:MP-VENUS (Bhatia et al., 2016). This fragment was inserted upstream of a 9X alanine linker followed by single YPET (Nguyen and Daugherty, 2005) or two tandem copies of YPET and the OCS terminator. DOF5.8 and TMO6 or TMO5 genomic coding sequences were then amplified using specific primers (Table S1) and cloned into a BamHI site, fused in-frame with the YPET tag to generate UBQ10p:lox spacer lox:GENE-YPET or UBQ10p:lox spacer lox:GENE-2xYPET. Finally, UBQ10p:lox spacer lox:GENE-2xYPET, and CLV3p:CRE-GR spacer lox were combined into the transfer DNA vector BGW (Karimi et al., 2002) using Gateway technology (Invitrogen). These constructs were then transformed into Agrobacterium strain C58C1 via electroporation and subsequently introduced into Arabidopsis plants using the floral dipping method (Clough and Bent, 1998).

Genomic sequences of DOF5.8, TMO6, TMO5, T5L1, T5L2 or T5L4 containing upstream and coding sequences were then amplified using specific primers (Table S1) and cloned into XhoI and BamHI sites, fused in-frame with a 9x alanine linker followed by YPET or 2xYPET tag to generate DOF5.8-YPET, TMO6-2YPET, TMO5-2YPET, T5L1::T5L1-2YPET, T5L2::T5L2-2YPET or T5L4::T5L4-2YPET translational reporter genes. Translational reporter genes were transferred as NotI restriction fragments to T-DNA vector pMLBART (Gleave, 1992).

### Chemical treatments

To induce DOF5.8/TMO5/TMO6 sectors in the *mp* mutant meristem, *mp* mutants harboring the respective sectoring constructs (CLV3p::CRE-GR + UBQ10p::lox spacer lox::GENE-2YPET) were germinated and grown on 1X MS agar medium until the emergence of leafless or flowerless dome structures. A 10–20 μL aliquot of 10 μM dexamethasone (DEX) in sterile water (prepared from a 10 mM stock dissolved in absolute ethanol) was directly applied to the dome meristem. Imaging was performed using confocal or brightfield microscopy at different time intervals, starting from 0 day and continuing up to 6–14 days. For the induction of same constructs in the wild type vegetative meristems, seeds were germinated on MS medium containing 10 μM DEX. The vegetative meristem was dissected 3 days after stratification (das), following the method described previously (Caggiano et al., 2021). Seedlings displaying sector formation were transferred to MS medium without DEX, and the meristem was imaged until 5–6 das. For overexpression of TMO5 (UBQ10p::lox spacer lox::TMO5-2YPET) in *mp* mutants, seedlings having dome meristems grown on MS medium were transferred to medium containing 10 μM DEX for continuous induction. Confocal or brightfield imaging was performed over a 6–14 day period. For overexpression in wild type vegetative meristem, seedlings were germinated and grown on MS medium containing 10 μM DEX. Vegetative meristems were dissected at 3das and imaged until 5 das. In all experiments, *mp* mutants were imaged before and after induction.

For N-1-naphthylphthalamic acid (NPA) treatment on the *mp* mutant (to inhibit polar auxin transport), the dome meristem harbouring the TMO5 sectoring construct (CLV3p::CRE-GR + UBQ10p::lox spacer lox::TMO5-2YPET) was pre-treated with 10 μM NPA for 2 days prior to DEX induction to eliminate residual auxin transport. After a single 10 μM DEX treatment, the mutants were continuously grown in the presence of 10 μM NPA or a DMSO mock treatment. Imaging was performed before and after chemical treatments and continued until 10 days post DEX treatment.

For NPA treatment in wild type and *tmo5/t5l* single and higher-order mutants, seedlings were grown from germination on MS medium containing 1 μM NPA or a DMSO mock treatment. Vegetative and inflorescence meristems were imaged using brightfield microscopy.

For auxinole treatment of the *mp* mutant (to inhibit auxin signalling), the dome meristem harboring the TMO5 sectoring construct was pre-treated with 100 μM auxinole or a DMSO mock for 2 days to eliminate residual auxin signaling. Following a single 10 μM DEX treatment on the dome meristem, the mutants were continuously grown in either 100 μM auxinole or a mock treatment. Imaging was performed before and after chemical treatments and continued until 10 days post-DEX treatment.

For organ outgrowth in the *mp* mutant, cytokinin (10 μM trans-zeatin, TZ) was applied to the *mp* vegetative meristem, while both auxin (5 mM IAA) and cytokinin (1 mM TZ) were applied to young pin-like inflorescence meristems, with DMSO/water as the mock control. Observations were recorded 7 to 14 days after treatment.

For auxin and cytokinin treatment on *tmo5* quadruple mutants (*tmo5, t5l1, t5l2*, and *t5l3*), a single application of 10 μM trans-zeatin (TZ) and 1 μM naphthaleneacetic acid (NAA) was administered directly to the shoot tip of 5-day-old seedlings. Treatments were applied either individually or in combination, with DMSO/water as the mock control. Additionally, 1 μM NAA or 1 μM or 0.5 μM TZ was included in the growth medium under some conditions. Observations were taken 10-20 days after treatment.

### Inflorescence Floral quantitation

To quantify the number of flowers in the inflorescence meristem in wild type and *tmo5* mutants, primary inflorescences were examined at the stage when the first flower bud had opened. The total number of flower buds within the inflorescence meristem was counted, and the average number of flower buds per inflorescence was calculated.

### Phyllotaxy measurement

To measure the phyllotactic pattern in wild type and *tmo5* mutants, the divergent angles between successive siliques in the mature inflorescence were assessed. The angles between each pair of siliques were recorded and averaged for comparison.

### Confocal live imaging

Wild type vegetative and inflorescence meristems were dissected for imaging as previously described (Caggiano et al., 2021; Heisler et al., 2005). Confocal live imaging was performed using a Leica TCS-SP5 upright laser scanning confocal microscope with hybrid detectors (HyDs). Imaging utilized a 25X water objective (N.A. 0.95) or a 63X objective (N.A. 1.20) and a pixel format of either 512×512 or 1024×1024 for enhanced resolution. The system was configured for bidirectional scanning at speeds of 400 Hz or 200 Hz, with line averaging set to 2 or 3, producing optical sections with a thickness of 1 μm.

The laser settings for GFP and YPet/VENUS, as described in previous studies (Kareem et al., 2022), were applied. CFP was excited using 456 nm laser and emission window was set to 465–500nm using argon laser. The pinhole was adjusted based on fluorescence brightness to avoid signal saturation or bleaching. A smart gain setting of 100% was employed, and sequential scan mode was utilized to switch between frames for imaging CFP/GFP and YPet simultaneously.

### Image analysis and data processing

Images were analyzed using Imaris 9.1.2 (Bitplane) or ImageJ (FIJI, https://fiji.sc). They were annotated and arranged in Adobe Photoshop (2020) and Affinity Designer 1.10.8. PIN1 polarity assessments were performed by examining arcs of fluorescent signal that extend beyond cell junctions and around cell corners, as previously described (Kareem et al., 2022).

**Fig. S1.**
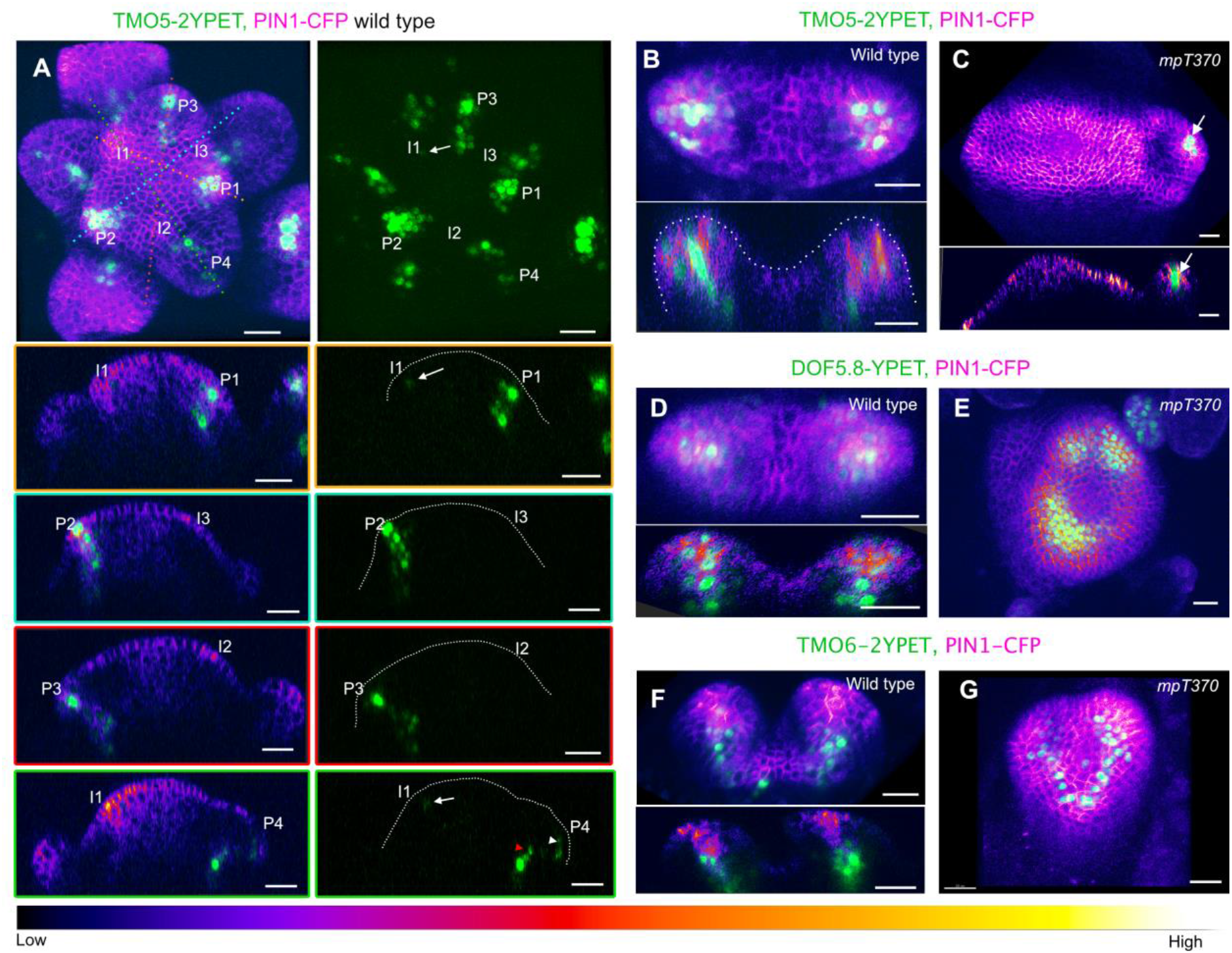
Expression pattern of TMO5, DOF5.8, and TMO6 in wild type and *mp* mutant meristems. (A) TMO5::TMO5-2YPET expression (green) in wild type inflorescence meristem partially overlapping with PIN1::PIN1-CFP (magenta) at various organ initiation stages. Incipient primordia (i3, i2, i1) and developing primordia (p1, p2, p3, p4) are labeled. Right panels show the green channel from the merged images. (B) TMO5::TMO5-2YPET expression overlapping with PIN1::PIN1-CFP in wild type vegetative meristem. Lower panel shows longitudinal optical section (C) TMO5::TMO5-2YPET expression (arrow) in the rarely formed leaves of the *mp-*T370 mutant dome meristem. Lower panel shows longitudinal optical section (D) DOF5.8::DOF5.8-YPET expression (green) wild type vegetative meristem. Lower panel shows longitudinal optical section (E) DOF5.8::DOF5.8-YPET expression in *mp*-T370 mutant pin-like meristem, showing reduced expression at later stages. (F) TMO6::TMO6-2YPET expression (green) in wild type vegetative meristem. Lower panel shows longitudinal optical section (G) TMO6::TMO6-2YPET expression in mpT370 mutant pin-like meristem. Scale bar = 20 μm.

**Fig. S2.**
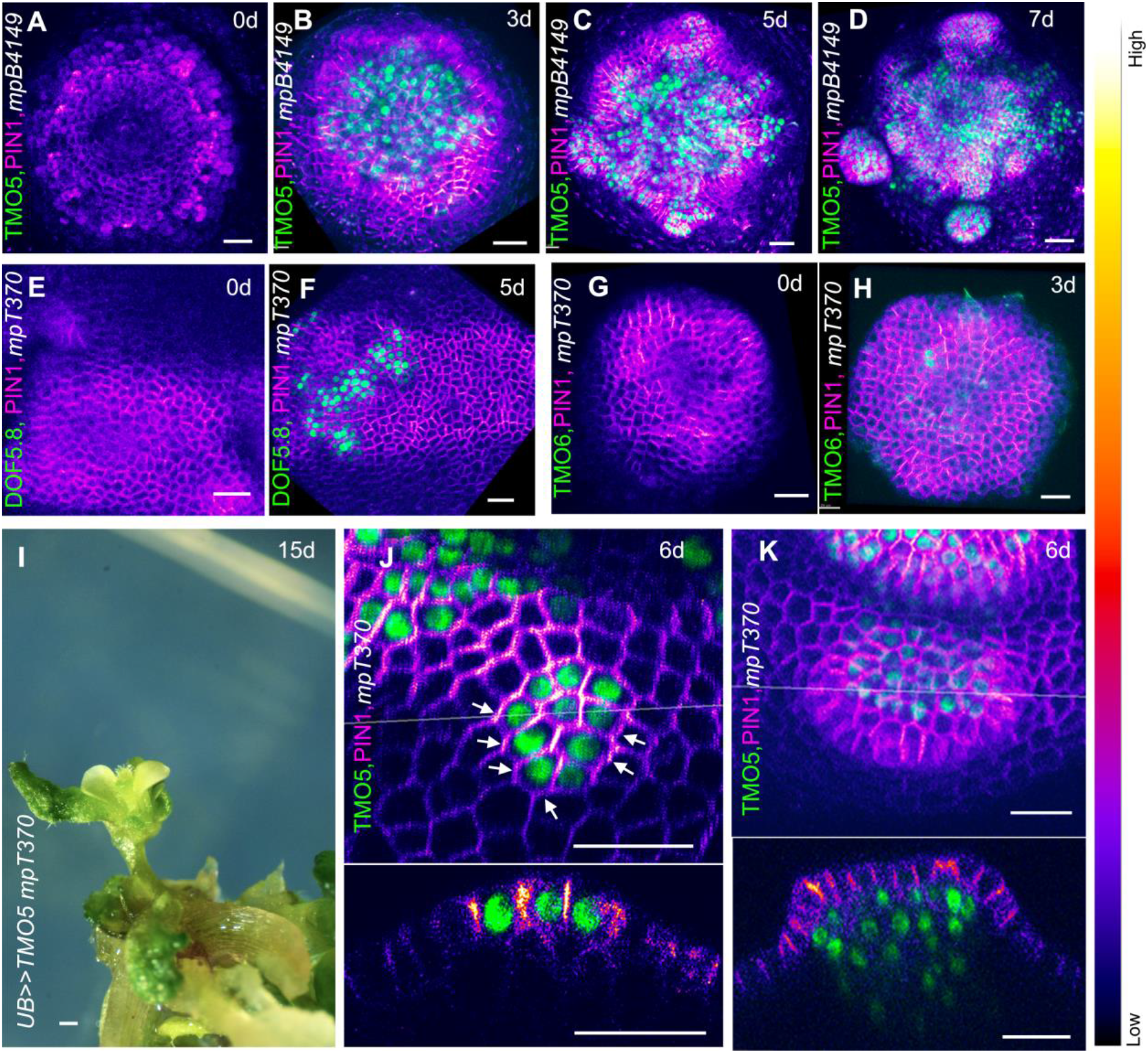
TMO5 rescues PIN1 polarization and organogenesis in *mp* mutant. (A-D) Activation of PIN1-GFP expression (magenta), convergence formation, and organ outgrowth in *mp* mutant shoot apical meristems following local activation of TMO5-2YPET clones (green). (E-H) No activation of PIN1-CFP expression or organ outgrowth in *mp* mutant after local expression of clones of DOF5.8-YPET (E,F) or TMO6-YPET (G,H). (I) Rescue of the *mp* mutant flower phenotype by transient ubiquitous expression of TMO5-2YPET with 8 days of Dex treatment, followed by 7 days without Dex. (J) PIN1 polarization towards TMO5-expressing clones in neighboring cells in *mp* mutant meristems. Lower panel shows the corresponding longitudinal optical section. Arrows denote PIN1 polarity. (K) Activation of PIN1 expression in epidermal cells in response to underlying sub-epidermal TMO5 clones. Lower panel shows the corresponding longitudinal optical section. Scale bar = 20 µm, except for (I) where it is 1 mm.

**Figure S3.**
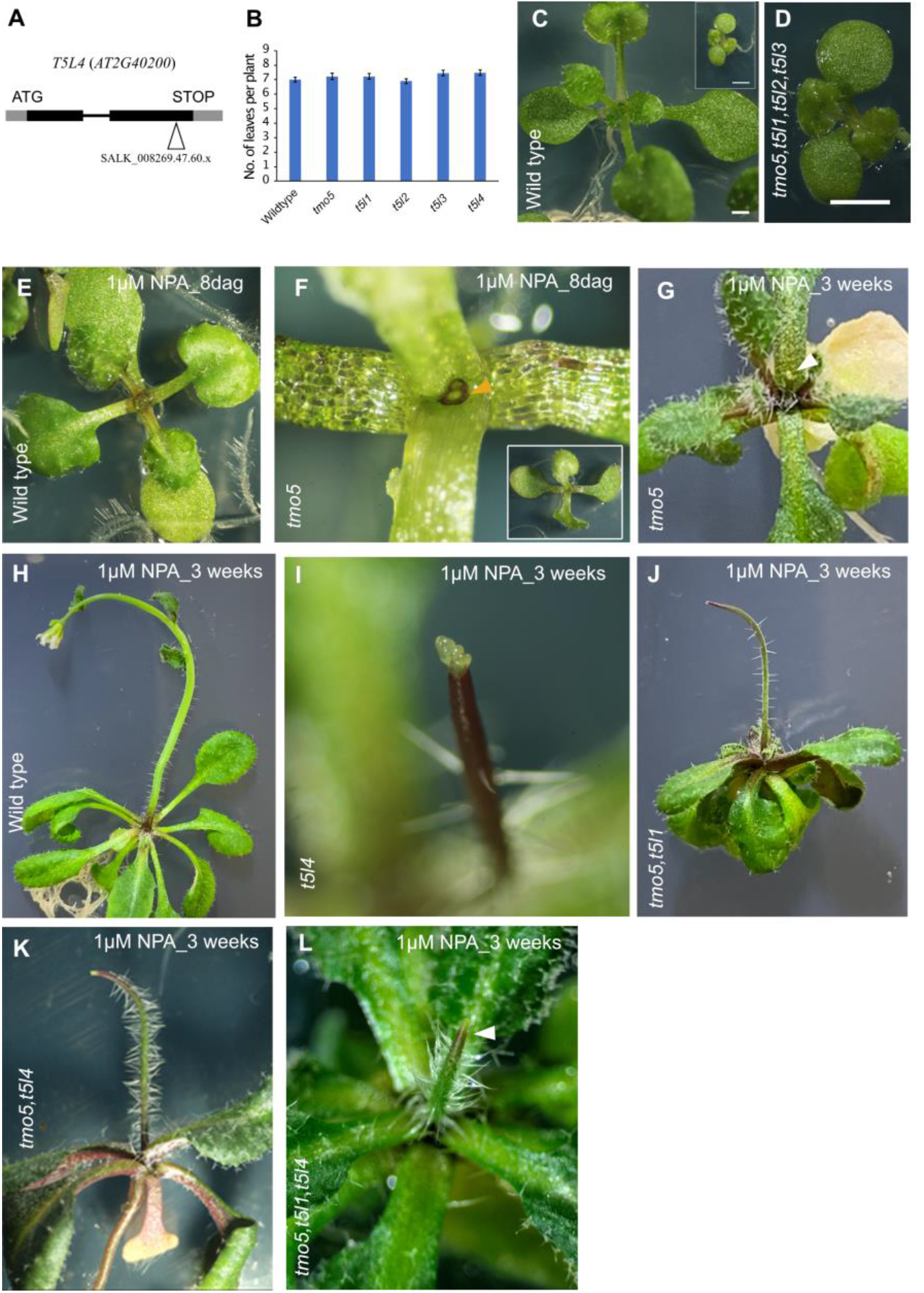
Phenotypic characterization of TMO5/T5L mutants. (A) Gene structure and location of T-DNA insertion of *t5l4* mutant (AT2G40200, SALK_008269.47.60, N508269) (B) Leaf formation in wild type and *tmo5/t5l* mutants. (n=31plants per genotype; mean±SEM). (C,D) Growth phenotypes of wild type and *tmo5,t5l1,t5l2,t5l3* quad mutant. Inset in (B) shows *tmo5* quad at the same scale as the wild type size. (E) Wild type seedling after 8 days of 1 μM NPA treatment. (F) Leafless dome structure in *tmo5* mutants upon 1 μM NPA treatment. Inset shows an overview image. (G) Pin-like inflorescence formation in *tmo5* mutants after 1 μM NPA treatment. (H) Wild type seedling after weeks of 1 μM NPA treatment. (I-L) Pin-like inflorescence produced by *tmo5,t5l1* double and *tmo5,t5l1,t5l4* triple mutants upon 1 μM NPA treatment. Yellow arrowhead in (F) indicates dome shaped meristem and white arrowheads in (G and L) indicate pin-like inflorescence. Scale bar: 1 mm.

**Fig. S4.**
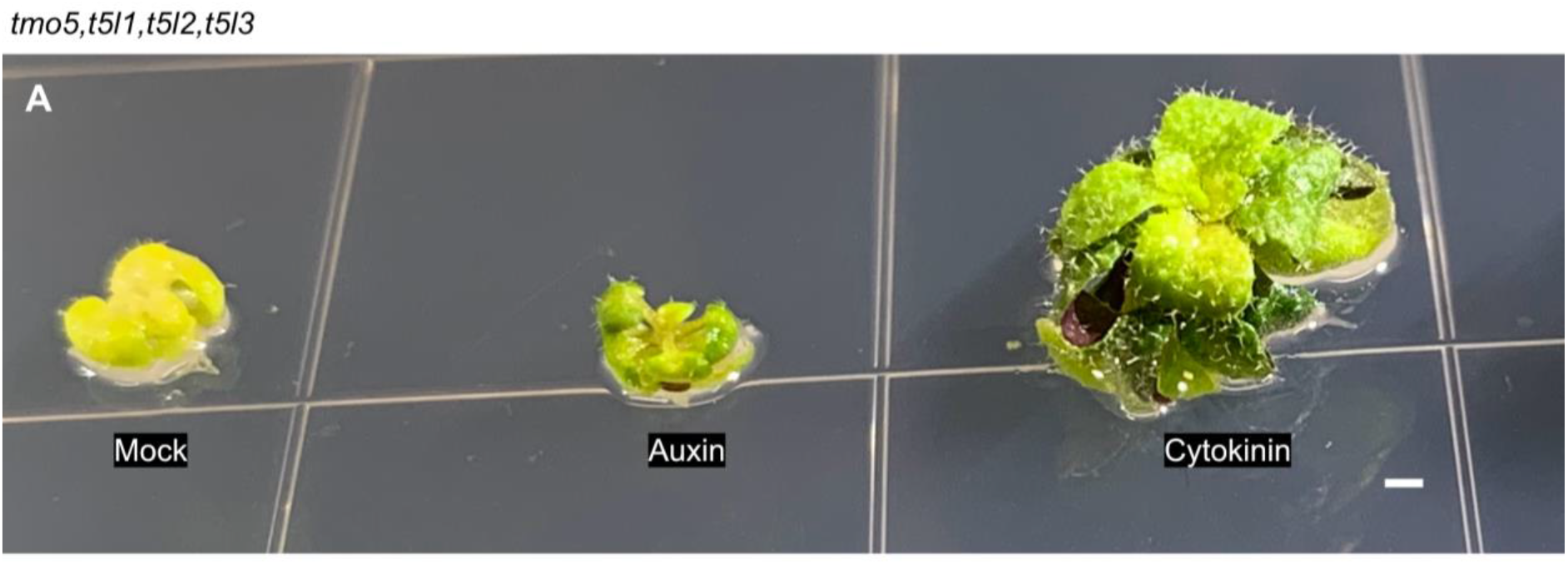
Cytokinin rescues leaf formation in *tmo5* quadruple mutant. (A) Leaf formation in the *tmo5,t5l1,t5l2,t5l3* quadruple mutant after treatment with 10 μM TZ (cytokinin), with no effect from 1 μM NAA (auxin) alone. Scale bar = 1 mm.

**Supplementary Table S1.**
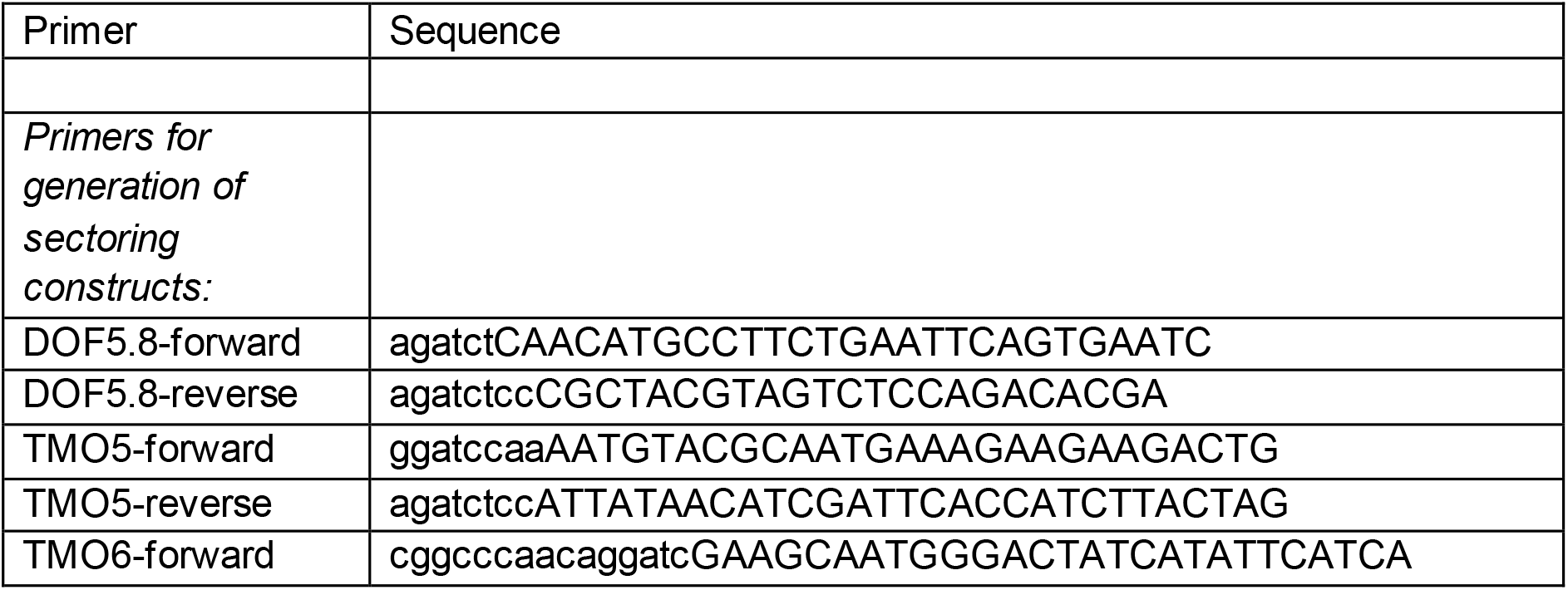
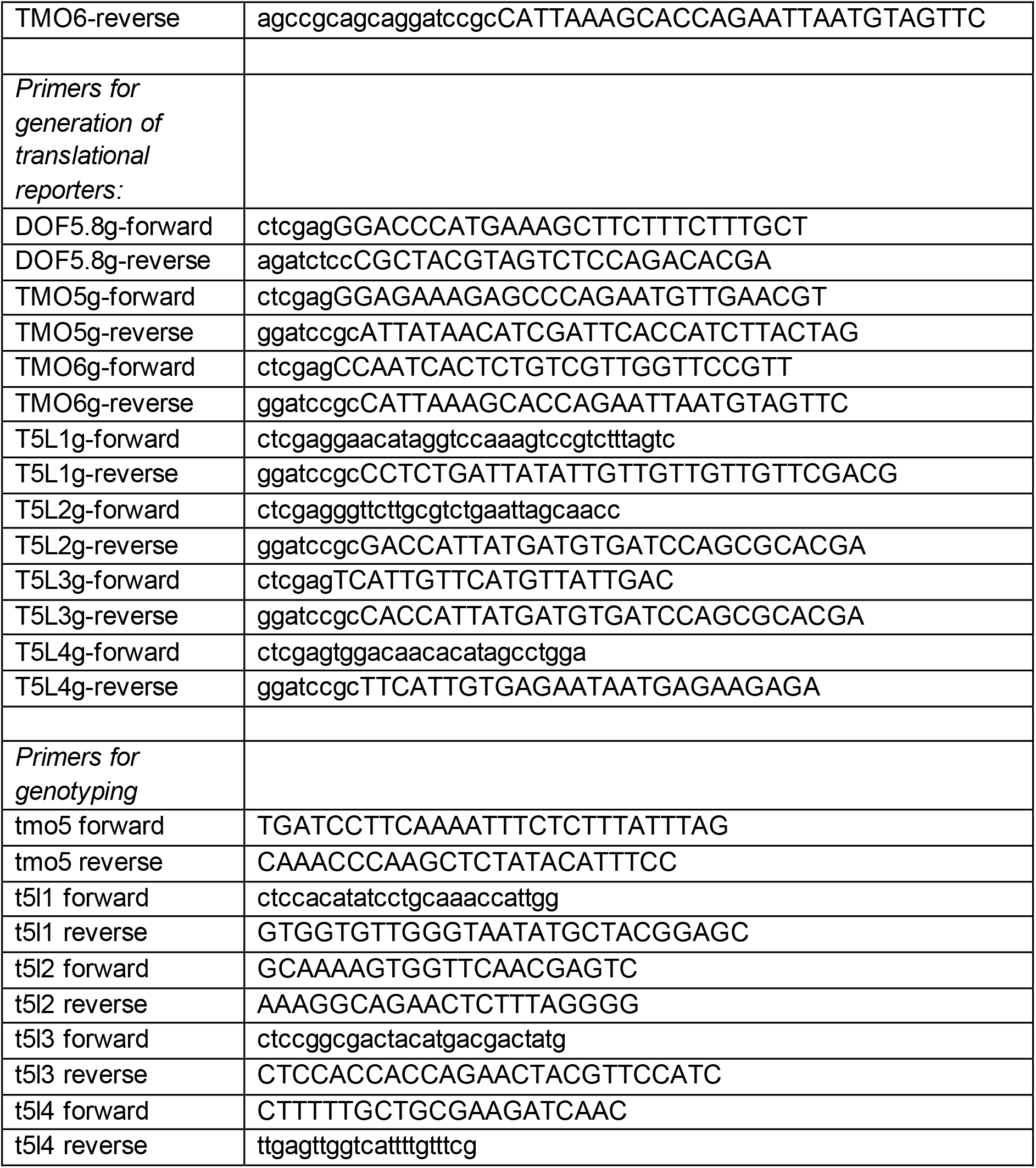

## Acknowledgment

This work was supported by the Australian Research Council Discovery Grant DP180101149 awarded to MGH.

## Author contribution

AK carried out the experiments and helped in their design and in the writing of the manuscript. CO helped provide experimental materials. MGH helped conceive the experiments and write the manuscript.

